# LiaR-dependent gene expression contributes to antimicrobial responses in group A *Streptococcus*

**DOI:** 10.1101/2024.04.04.588141

**Authors:** Luis Alberto Vega, Misu Sansón-Iglesias, Piyali Mukherjee, Kyle Buchan, Gretchen Morrison, Anne E. Hohlt, Anthony R. Flores

## Abstract

The ability to sense and respond to host defenses is essential for pathogen survival. Some mechanisms involve two-component systems (TCS) that respond to host molecules, such as antimicrobial peptides (AMPs) and activate specific gene regulatory pathways to aid in survival. Alongside TCSs, bacteria coordinate cell division proteins, chaperones, cell wall sortases and secretory translocons at discrete locations within the cytoplasmic membrane, referred to as functional membrane microdomains (FMMs). In Group A *Streptococcus* (GAS), the FMM or “ExPortal” coordinates protein secretion, cell wall synthesis and sensing of AMP-mediated cell envelope stress via the LiaFSR three-component system. Previously we showed GAS exposure to a subset of AMPs (α-defensins) activates the LiaFSR system by disrupting LiaF and LiaS co-localization in the ExPortal, leading to increased LiaR phosphorylation, expression of the transcriptional regulator SpxA2, and altered GAS virulence gene expression. The mechanisms by which LiaFSR integrates cell envelope stress with responses to AMP activity and virulence are not fully elucidated. Here, we show the LiaFSR regulon is comprised of genes encoding SpxA2 and three membrane-associated proteins: a PspC domain-containing protein (PCP), the lipoteichoic acid-modifying protein LafB and the membrane protein insertase YidC2. Our data show phosphorylated LiaR induces transcription of these genes via a conserved operator, whose disruption attenuates GAS virulence and increases susceptibility to AMPs in a manner primarily dependent on differential expression of SpxA2. Our work expands understanding of the LiaFSR regulatory network in GAS and identifies targets for further investigation of mechanisms of cell envelope stress tolerance contributing to GAS pathogenesis.

## Introduction

The bacterial response to human host molecules is critical in the development of disease. Antimicrobial peptides (AMPs), produced by immune and epithelial cells of skin and mucosal surfaces, are part of the first line of defense against bacterial infection (Cole and Nizet 2016). Through multiple mechanisms, AMPs induce cell envelope stress that may lead to bacterial killing and clearance (Melo, Ferre, and Castanho 2009; Pachón-Ibáñez et al. 2017). In response, bacterial pathogens have evolved to sense, inactivate and counter AMP activity to survive. Resistance to the killing activity of AMPs is typically associated with changes in the bacterial cell envelope and involves membrane proteins that mediate host-pathogen interaction (Assoni et al. 2020). Given their critical role in response to infection, bacteria have also developed gene regulatory systems to respond to the AMP-induced disturbances in essential cell processes. For example, bacterial two component systems (TCSs) (Stock, Robinson, and Goudreau 2000) activate gene regulatory pathways that contribute to AMP resistance and may further alter bacterial gene expression to respond to host-induced stress (Cho et al. 2023; Cheung et al. 2021; Chen et al. 2021; Chaili et al. 2016). The continued rise of antimicrobial resistance (Oppegaard et al. 2020; Fay et al. 2021; M. A. Sanson et al. 2019) and the potential role of AMPs as treatment alternatives (Lopes et al. 2022) necessitates a better understanding of bacterial response mechanisms to AMP activity.

The LiaFSR system responds to cell envelope stress, including that induced by AMPs. Conserved among Gram-positive bacteria, this TCS consists of a histidine kinase (HK) (LiaS), a response regulator (RR) (LiaR) and an accessory membrane protein (LiaF). LiaS is unique in that it is intramembrane-sensing and does not possess a large extracellular sensing domain typical of other HKs (Mascher 2006). Previously, it was shown that LiaS and LiaF localize at discrete locations within the cytoplasmic membrane of bacteria called functional membrane microdomains (FMMs) (García-Fernández et al. 2017; LaRocca et al. 2013; López and Kolter 2010) – referred to as the ExPortal in GAS (J. Rosch and Caparon 2004). Human α-defensins such as human neutrophil peptide 1 (hNP-1), highly expressed in the azurophilic granules of neutrophils (Ganz et al. 1985; Lehrer 1997), specifically disrupt the ExPortal, inhibiting protein secretion and processing (J. W. Rosch and Caparon 2005; Vega and Caparon 2012). GAS, like other streptococci, has in turn evolved mechanisms to respond to human AMP activity (Majchrzykiewicz, Kuipers, and Bijlsma 2010; Velarde, Ashbaugh, and Wessels 2014).

Activation of the LiaFSR system is known to have diverse effects among Gram-positive bacteria. In *Streptococcus mutans*, the LiaFSR system is one of the regulatory systems contributing to maintenance of cell wall and membrane homeostasis in response to oxidative stress by controlling expression of the transcriptional regulator SpxA2 (Ganguly et al. 2020; Baker et al. 2020). The *Streptococcus agalactiae* LiaFSR system was found to regulate expression of genes involved in cell wall synthesis, pili formation and cell membrane modification (Klinzing et al. 2013). In the same study, Klinzing et al. also showed that deletion of LiaR resulted in increased susceptibility to cell wall-active antibiotics (vancomycin and bacitracin) and AMPs (polymixin B, colistin, and nisin) and significant attenuation of virulence in mouse models of *S. agalactiae* sepsis and pneumonia. Recently, there has been increased interest in LiaFSR regulatory mechanisms, since mutations in this system have been associated with reduced daptomycin susceptibility and increased sensitivity to glycopeptides in clinically relevant *Enterococcus* species (Ota et al. 2021; Prater et al. 2021; Gargis et al. 2021; Zeng et al. 2022; Tymoszewska, Szylińska, and Aleksandrzak-Piekarczyk 2023; Axell-House et al. 2024).

In previous studies we showed that GAS exposure to hNP-1 and cell membrane-active antimicrobials disrupt LiaF and LiaS co-localization in the ExPortal leading to increased LiaR phosphorylation and increased transcription of *spxA2* (Lin et al. 2020; M. A. Sanson et al. 2021). LiaR-related changes in *spxA2* transcript levels were associated with altered GAS virulence gene expression (e.g. *speB, mac, grab*) and changes in phosphorylation of the virulence regulator CovR (M. A. Sanson et al. 2021). However, the mechanisms by which LiaFSR integrates cell envelope stress with responses to AMP activity and virulence are not fully elucidated. In the current study we demonstrate that LiaR phosphorylation induces expression of the LiaFSR-specific regulon, limited to genes encoding SpxA2 and three membrane-associated proteins. We show that phosphorylated LiaR influences transcription via a conserved operator sequence. Disruption of LiaR gene regulation results in attenuated fitness in an *ex vivo* model of infection and increased susceptibility to AMPs that is primarily dependent on differential expression of SpxA2. Our work expands understanding of the LiaFSR regulatory network in GAS and identifies targets for further investigation of mechanisms of cell envelope stress tolerance contributing to GAS pathogenesis.

## Results

### Phosphorylation of LiaR is required for differential expression of the LiaFSR regulon

Our previous studies revealed an inverse correlation in global gene expression between strains lacking either LiaF (Δ*liaF*, high LiaR phosphorylation) or LiaR (Δ*liaR*) compared to the serotype *emm3* isogenic parent MGAS10870 (M. A. Sanson et al. 2021). Thus, we hypothesized that LiaR phosphorylation is required for LiaFSR-associated changes in global gene expression in response to cell envelope-targeting antimicrobials. The LiaFSFR system is activated under conditions of cell envelope stress (e.g., antimicrobials and AMPs) that may also influence other gene regulatory systems and thus result in varied phenotypes (M. A. Sanson et al. 2021; Ganguly et al. 2020). To test our hypothesis and avoid the potentially confounding effects of antimicrobial exposure, we measured global gene expression using isogenic, single amino acid replacements in LiaS with differing levels of LiaR phosphorylation. Based on previous studies of HKs identifying conserved residues essential for cognate RR phosphorylation and dephosphorylation (Huynh, Noriega, and Stewart 2010) (**Figure 1A**), we generated an isogenic mutant in LiaS (Q146A) that lacks the ability to dephosphorylate LiaR. As predicted, we observed significantly increased LiaR phosphorylation (**Figure 1A**) and transcript levels of *spxA2* (**Figure 1B**), a gene previously shown to be directly regulated by LiaR (M. A. Sanson et al. 2021), in the Q146A mutant compared to the parental strain. Importantly, both LiaR phosphorylation and *spxA2* transcription were similar to those observed following LiaFSR activation by bacitracin in the isogenic MGAS10870 parent (**Figure 1A**, **B**).

**Figure 1.**
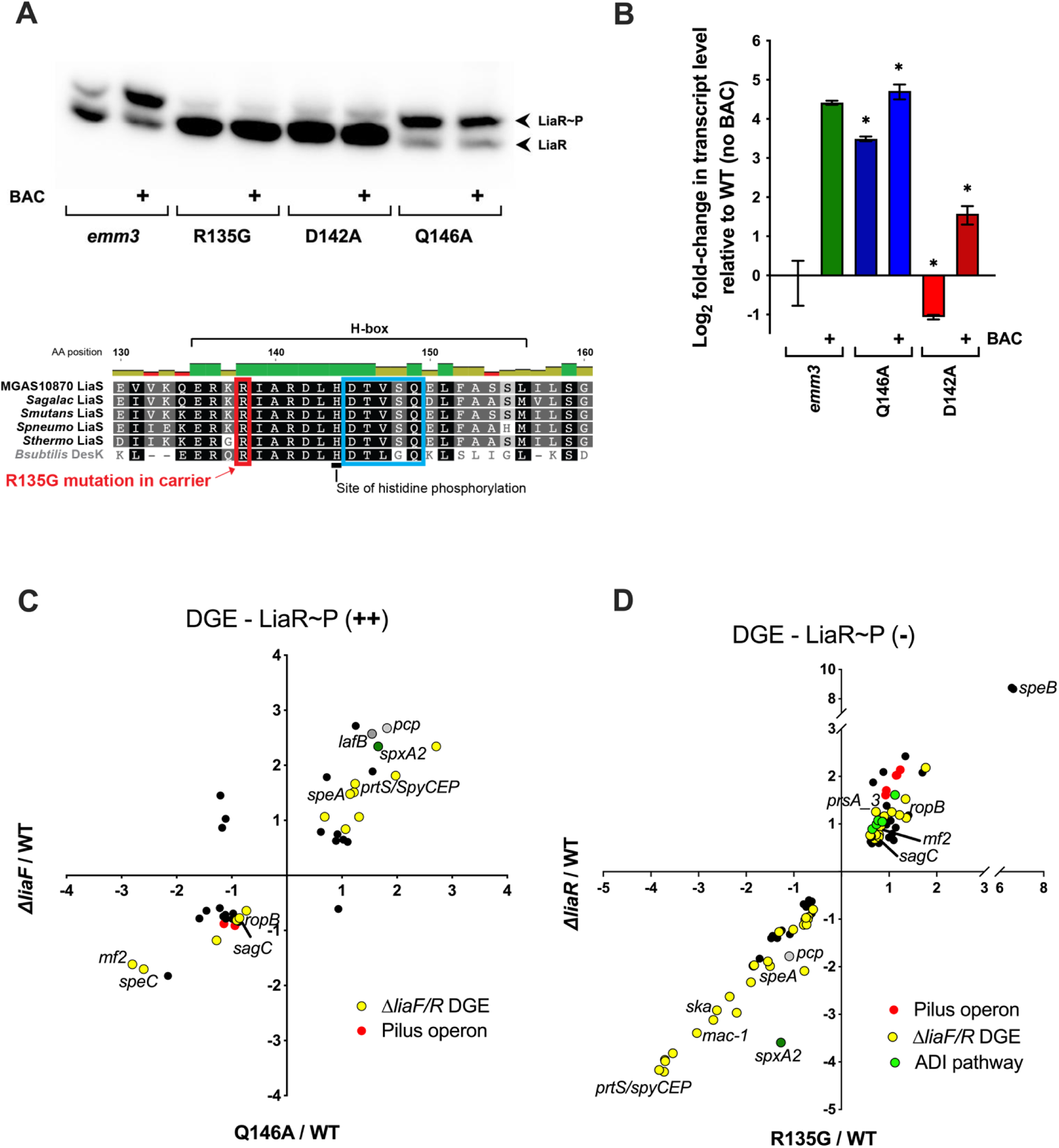
Opposing patterns of of LiaR phosphorylation result in differential transcription of spxA2 and correlate with ΔliaF and ΔliaR transcriptomes. (**A**) Mutagenesis of amino acid residues of the LiaS kinase involved in phosphorylation/dephosphorylation of LiaR protein (Q146, D142, R135) results in altered levels of phosphorylated LiaR in the presence/absence of exposure to bacitracin (BAC). Substitutions for Ala residues (Q146A, D142A), and the previously isolated R135G mutation associated with altered LiaFSR activity are shown. (**B**) Measurement of *spxA2* transcript levels by quantitative real-time PCR (qRT-PCR) in WT MGAS10870, Q146A and D142A isogenic LiaS mutants exposed to BAC confirms *spxA2* transcript levels correlate with LiaR phosphorylation. (**C**) Correlation plot of 41 significantly (*P* < 0.05; Bonferroni correction) differentially expressed genes (≥1.5-fold relative to WT MGAS10870) shared between the Q146A (x-axis) and the Δ*liaF* (y-axis) transcriptome. (**D**) Correlation plot of 86 significantly (*P* < 0.05, Bonferroni correction) differentially expressed genes (≥1.5-fold relative to WT MGAS10870) shared between the emergent R135G (x-axis) and the Δ*liaR* (y-axis) transcriptome. Log_2_ values are plotted, colors correspond to operons or individual genes whose differential expression was inversely correlated in Δ*liaF* and Δ*liaR* transcriptomes (Δ*liaF/R* DGE, previously published in (Sanson et al. 2021)) Names of virulence genes of interest are listed. Bacitracin exposure = 1µg/ml, 1h, mid-log phase, 37°C. Transcripts assessed by qRT-PCR analysis were measured in technical triplicate of biological quadruplicates, with significance determined by a t-test (*,*P* < 0.05; **, *P* < 0.01). Data is presented as the mean transcript level with standard error of the mean (SEM). Transcriptomes were analyzed by RNA-seq in biological triplicate and DGE determined for core genome only, excluding phage and other mobile elements.

We next sought to identify a mutation in LiaS that would eliminate cognate RR phosphorylation. Previously, we described an arginine (R) to glycine (G) mutation at position 135 in LiaS (**Figure 1A**) that contributed to altered virulence and a carrier phenotype in a GAS clinical isolate (Flores et al. 2015) (R135G, carrier mutation). Given that our previous studies showed phenotypic and gene regulatory similarities between the R135G mutant and a Δ*liaR* strain (Flores et al. 2017; M. A. Sanson et al. 2021), we hypothesized that the R135G carrier mutation significantly reduced LiaR phosphorylation in GAS. To test this hypothesis, we also generated a single amino acid replacement in LiaS (aspartate to alanine at position 142, D142A) predicted to lack the ability to phosphorylate LiaR (Trajtenberg et al. 2010; Huynh, Noriega, and Stewart 2010) as a control and compared the mutants and parental strain LiaR phosphorylation and *spxA2* transcript levels. We observed nearly absent LiaR phosphorylation (**Figure 1A**) and reduced levels of *spxA2* transcript (**Figure 1B**) relative to the MGAS10870 parental strain in the D142A mutant and the R135G strain (Flores et al. 2017), independent of LiaFSR activation by bacitracin. These data confirm decreased LiaR phosphorylation as the mechanism for the gene regulatory and phenotypic changes associated with the R135G carrier mutation. Further, our data support the LiaS-Q146A (LiaR hyperphosphorylation) and LiaS-R135G (unphosphorylated LiaR) mutants in assessing the downstream regulatory effects associated with LiaR phosphorylation status.

We next used RNA sequencing (RNA-seq) to compare transcriptomes between the parental (MGAS10870) and isogenic LiaS mutants. Analysis of differential gene expression relative to the parental strain (≥1.5-fold change, *P* < 0.05 adjusted for multiple comparisons, core genome) showed up-regulation of 136 genes and down regulation of 98 genes in the Q146A mutant (Supplemental Table S1). Comparison to the published Δ*liaF* transcriptome revealed direct correlation of 41 genes (16.8% of Q146A transcriptome) differentially expressed under high levels of LiaR phosphorylation (**Figure 1C**). The comparison demonstrated multiple genes whose differential expression was shown to be inversely correlated in Δ*liaF* and Δ*liaR* mutants (Δ*liaF/R* DGE; yellow dots), including genes encoding known virulence determinants (*prtS/spyCEP*, *sdn*, *speA*, *speC*, *mf2*, *sagC*, *sdaB*) and members of the pilus operon (*srtB*, *tee3*). Also differentially expressed were multiple genes encoding transcriptional regulators known to be associated with virulence (*ropB*, *rgg3*, *msmR*) in addition to *spxA2* – a gene known to be directly regulated by LiaR (M. A. Sanson et al. 2021). Other potential contributors to GAS fitness and virulence of interest are ABC transporters (*braB/brnQ*, *dppE*), and the D-alanyl carrier protein involved in teichoic acid D-alanylation (*dltC*).

Given our previously published data indicating the R135G carrier mutation in LiaS contributes to GAS virulence and fitness phenotypes, we also assessed the transcriptome of the isogenic R135G mutant relative to the parental strain. Analysis of differential gene expression relative to the parental strain showed up-regulation of 68 genes and down regulation of 47 genes in the R135G mutant (Supplemental Table S2). Comparison to the published ΔliaR transcriptome reveals direct correlation of 86 (74.8% of R135G transcriptome) genes differentially expressed in the absence of LiaR phosphorylation (**Figure 1D**). Expression of *spxA2* is downregulated under both reduced phosphorylation of LiaR (LiaS-R135G) and in a LiaR deletion background, as are genes shown to be inversely correlated in Δ*liaF* and Δ*liaR* mutants (Δ*liaF/R* DGE), including genes encoding multiple virulence factors (*prtS/SpyCEP*, *hasABC*, *mac-1*, *ska*, *speA*, *nga*, *slo*). Conversely, the *emm3* pilus operon is upregulated (*cbp*, *tee3*, *srtAB*), as are *grab*, *mf2*, *sagC* and *speB* virulence factors, alongside virulence associated regulator *ropB* and the branched chain amino acid transporter *braB/brnQ*. Consistent with changes in LiaR phosphorylation status, comparison of DGE relative to MGAS10870 of the Q146A (increased LiaR phosphorylation) and R135G (decreased LiaR phosphorylation) transcriptomes revealed 30 inversely correlated transcripts, several of which were also inversely correlated in a comparison of Δ*liaF* and Δ*liaR* transcriptomes (M. A. Sanson et al. 2021) (e.g. *spxA2*, *ropB*, *prtS/spyCEP*, *speA3*, *tee3*) (Table 1, Supplemental Figure 1). Additionally, the operon encoding the arginine deiminase (ADI) pathway (*SpyM3_1192-96*), which contributes to GAS pathogenesis in mouse models of skin infection and resistance to killing by macrophages via iNOS inactivation (Hirose et al. 2021; Cusumano, Watson, and Caparon 2014), also displayed inversely correlated differential expression under distinct levels of LiaR phosphorylation (**Figure 1D** and Supplemental Figure 1; green dots). Altogether, these data support the distinct transcriptomic effects of LiaR phosphorylation and probable contribution to survival and virulence of GAS in response to cell envelope-targeting antimicrobials.

**Table 1.**
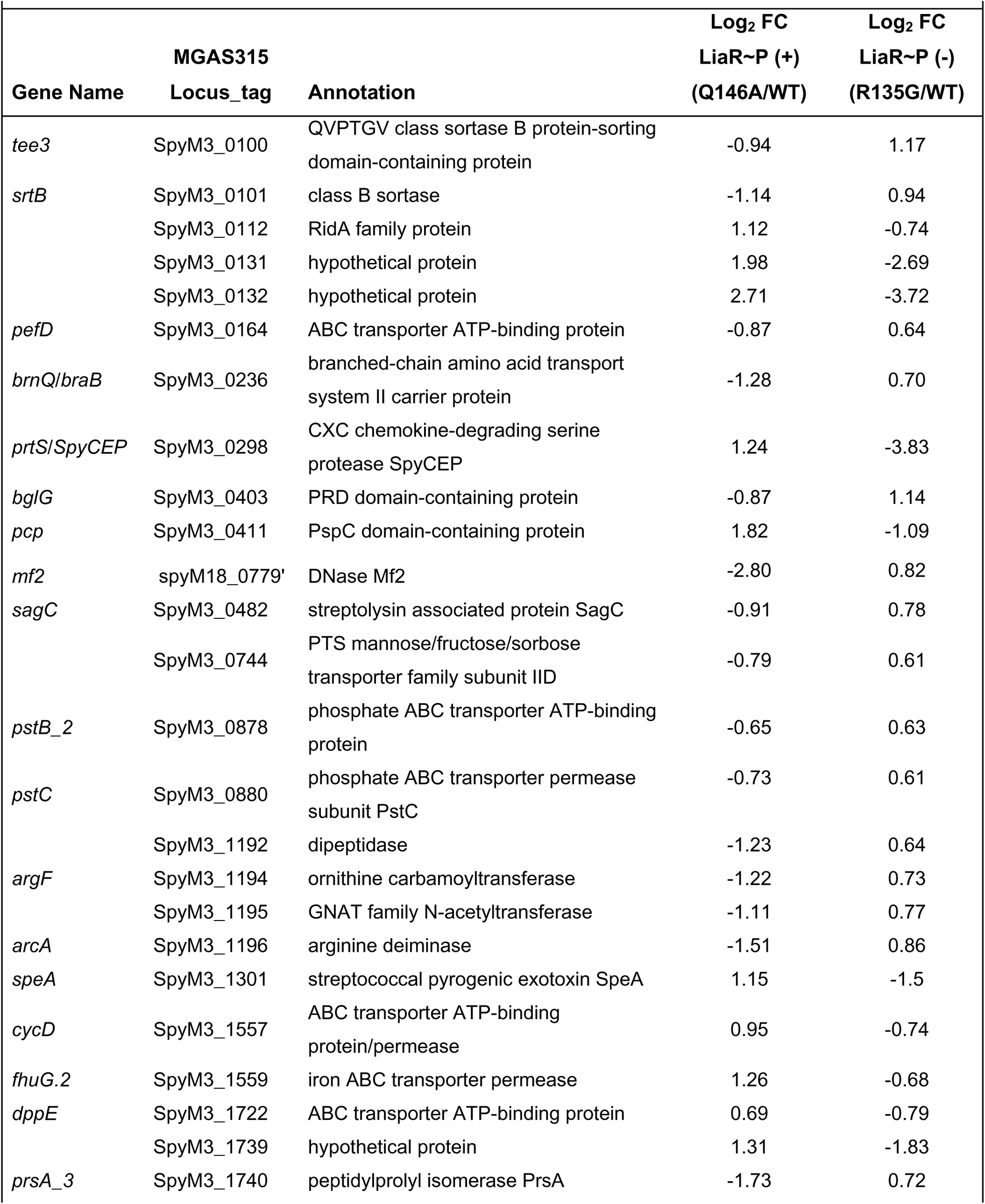
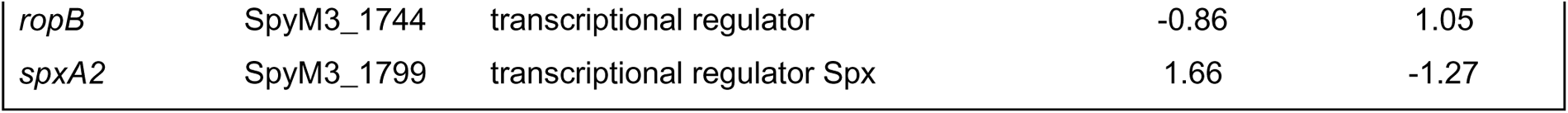
GAS differential gene expression associated with LiaR phosphorylation status.

### LiaR directly regulates a small subset of genes via a conserved operator

The transcriptomes of R135G and Q146A mutant strains combined with our previous work (M. A. Sanson et al. 2021) confirm direct regulation of *spxA2* by LiaR and suggest potential LiaR regulation of multiple additional targets including virulence determinants. We hypothesized that global gene regulation through phosphorylated LiaR may, in part, be expanded through direct regulation of *spxA2* and, thus, direct gene regulation of additional targets by LiaR more limited than predicted by our RNA-seq data. To test this hypothesis, we first defined global LiaR binding by measuring enrichment of GAS genomic sequences bound to phosphorylated LiaR using chromatin immunoprecipitation sequencing (ChIP-seq). In addition to the previously identified sequences in the *spxA2* promoter region (M. A. Sanson et al. 2021), ChIP-seq analysis identified enrichment of 3 gene promoter sequences in the Q146A strain relative to Δ*liaR* (negative control): SpyM3_0256 (*yidC2*), SpyM3_0363 (*lafB*) and SpyM3_0411 (*pcp*) (**Figure 2A**, Table 2). Analysis of Q146A and R135G transcriptomes showed inversely correlated differential expression of *pcp* relative to the isogenic MGAS10870 strain (**Figure 1C and D**, Supplemental Figure 1) in a manner resembling *spxA2* expression. Similarly, the *lafB* transcript was significantly increased in both the Δ*liaF* and Q146A transcriptomes (**Figure 1C**) but not in the Δ*liaR* and R135G backgrounds. Levels of the *yidC2* transcript were increased in the Δ*liaF* transcriptome (p=0.07, Bonferroni corrected) but not in the Q146A transcriptome. Using the MEME suite (Bailey et al. 2009), we next identified a single predicted consensus operator sequence (AnnAnnnnnCTnnnGnCnnATTTTT) in each of the promoters of the identified LiaR targets (Table 2). Analysis by qRT-PCR of *spxA2*, *lafB*, *pcp* and *yidC2* transcripts from MGAS10870 exposed to bacitracin (0.5 ug/mL) *in vitro* (rich medium, mid-exponential phase) revealed significantly increased levels of *spxA2*, *pcp*, and *lafB* transcripts relative to untreated culture (*P<*0.05; **Figure 2B**), further suggesting direct regulation by phosphorylated LiaR in response to cell envelope stress.

**Figure 2.**
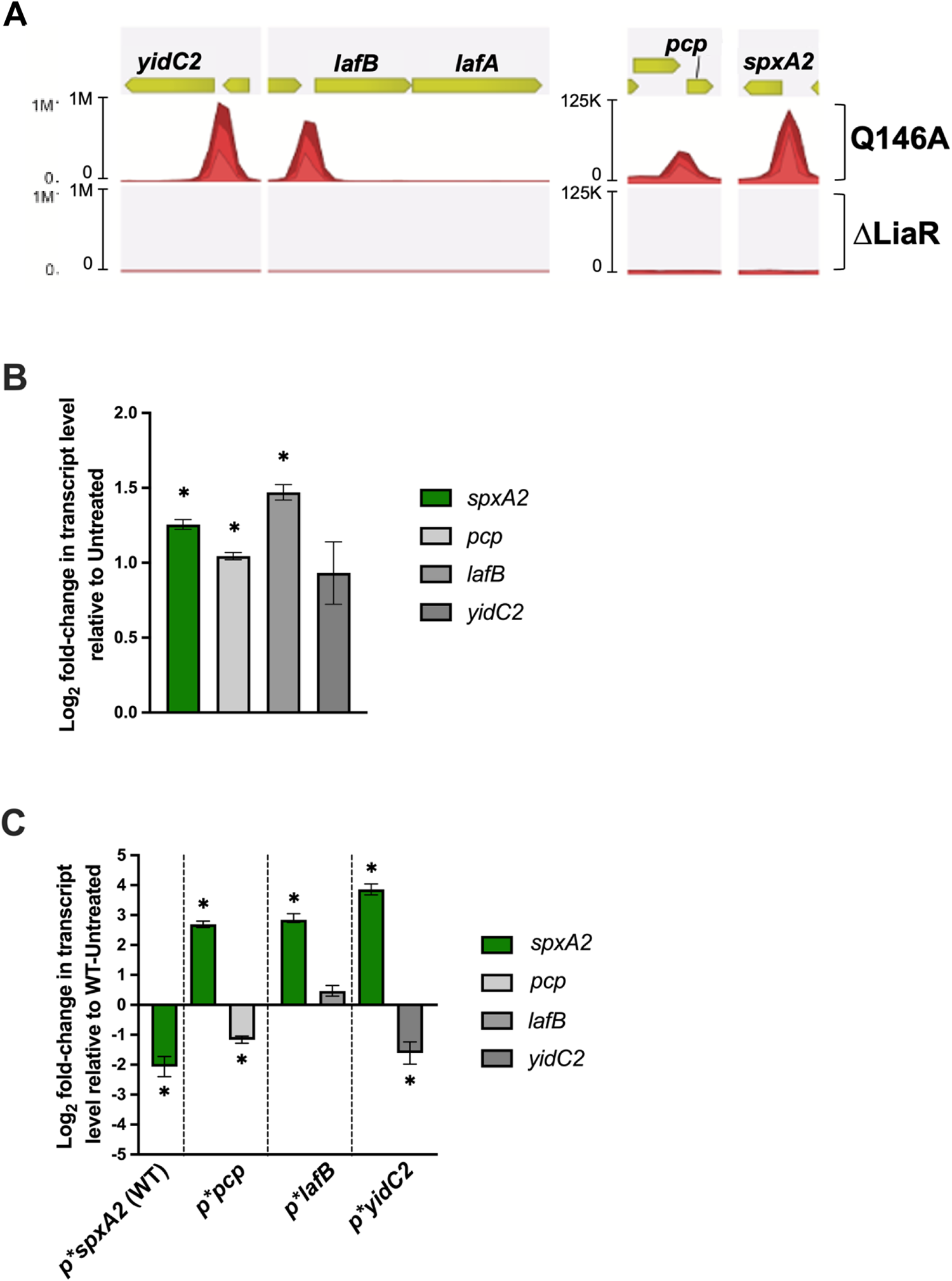
The regulon directly controlled by LiaR consists of *spxA2*, *yidC2*, *lafB*, and *pcp* transcripts. (**A**) Identified peaks and read mapping results of ChIP-seq analysis of the Q146A mutant relative to Δ*liaR* (negative control) reveal four transcripts directly regulated by phosphorylated LiaR. Quantitative real-time PCR (qRT-PCR) measurement of identified targets directly regulated by phosphorylated LiaR (*spxA2*, SpyM3_1799; *pcp*, SpyM3_0411; *lafB*, SpyM3_0363; *yidC2*, SpyM3_0256) in the bacitracin-exposed (1µg/ml, 1h, mid-log phase culture, 37°C) (**B**) wild-type (WT) MGAS10870 strain and (**C**) LiaR operator scramble mutants (p\**spxA2*, p\**pcp*, p\**lafB*, p\**yidC2*). Transcripts were measured in technical triplicate of biological quadruplicates, with significance determined by t-test (*, *P* < 0.05). Data is presented as the mean transcript level with standard error of the mean (SEM) relative to untreated WT MGAS10870 strain.

**Table 2.**
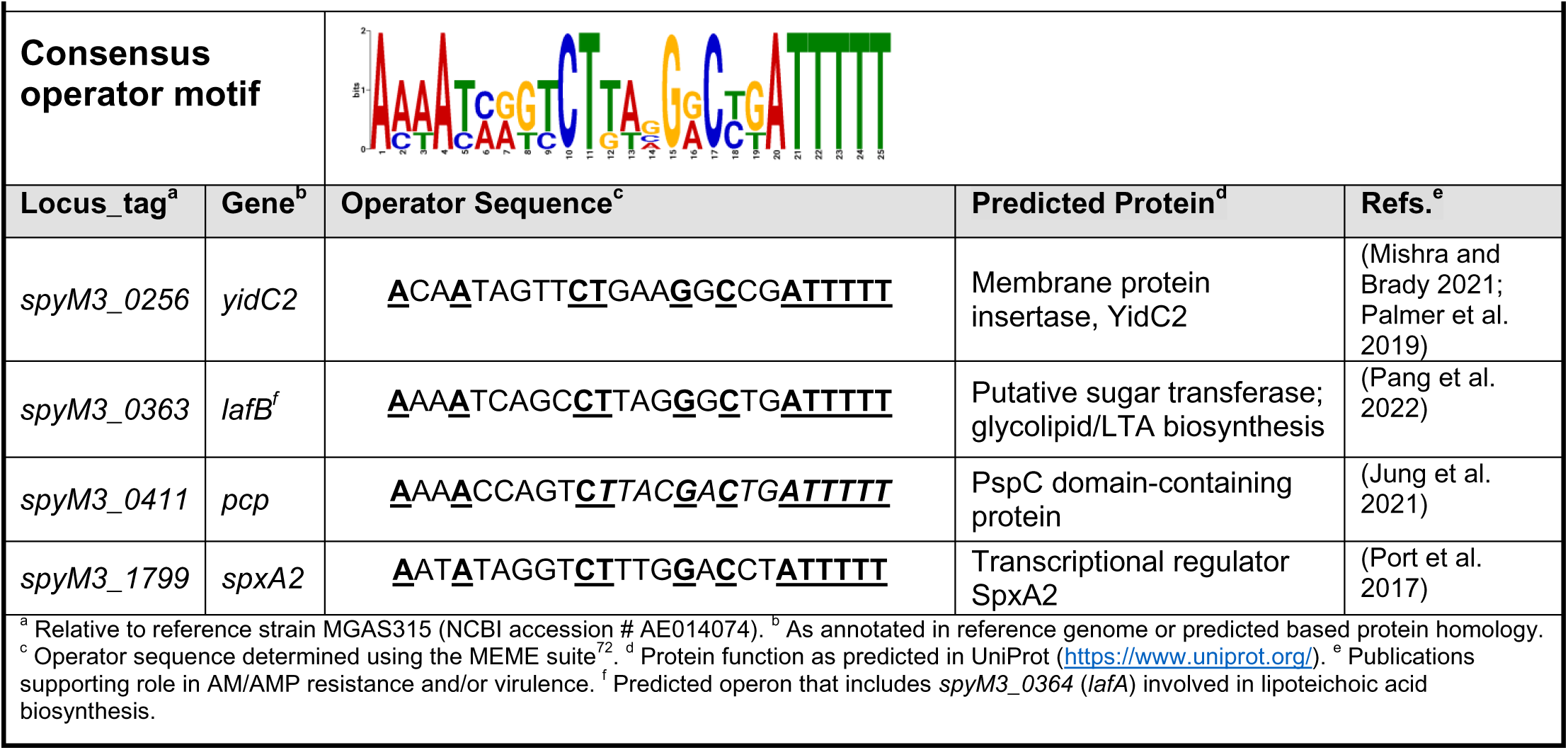
LiaR-regulated genes defined by ChIP-seq in *emm3* GAS.

Examination of 273 published, complete GAS genomes (NCBI) revealed the identified consensus operator sequence is highly conserved in GAS. The identified operator sequence was also highly similar to that described for the homologous genes shown to be regulated by LiaR in *S. mutans* (Shankar et al. 2015). Further, we did not identify additional predicted LiaR binding sites based on the consensus operator sequence in the reference MGAS10870 genome. We next hypothesized that the identified operator is necessary for LiaR-dependent gene regulation. We tested this hypothesis by generating independent isogenic operator “scramble” mutants of the identified LiaR targets in a wild-type strain background (p\**spxA2*, p\**pcp*, p\**yidC2*, p\**lafB*; Table 3). Mutagenesis of individual operator sequences resulted in significantly reduced gene transcript levels following bacitracin exposure relative to an untreated wild-type control, as determined by qRT-PCR analysis (*P*<0.05; **Figure 2C**).

**Table 3.**
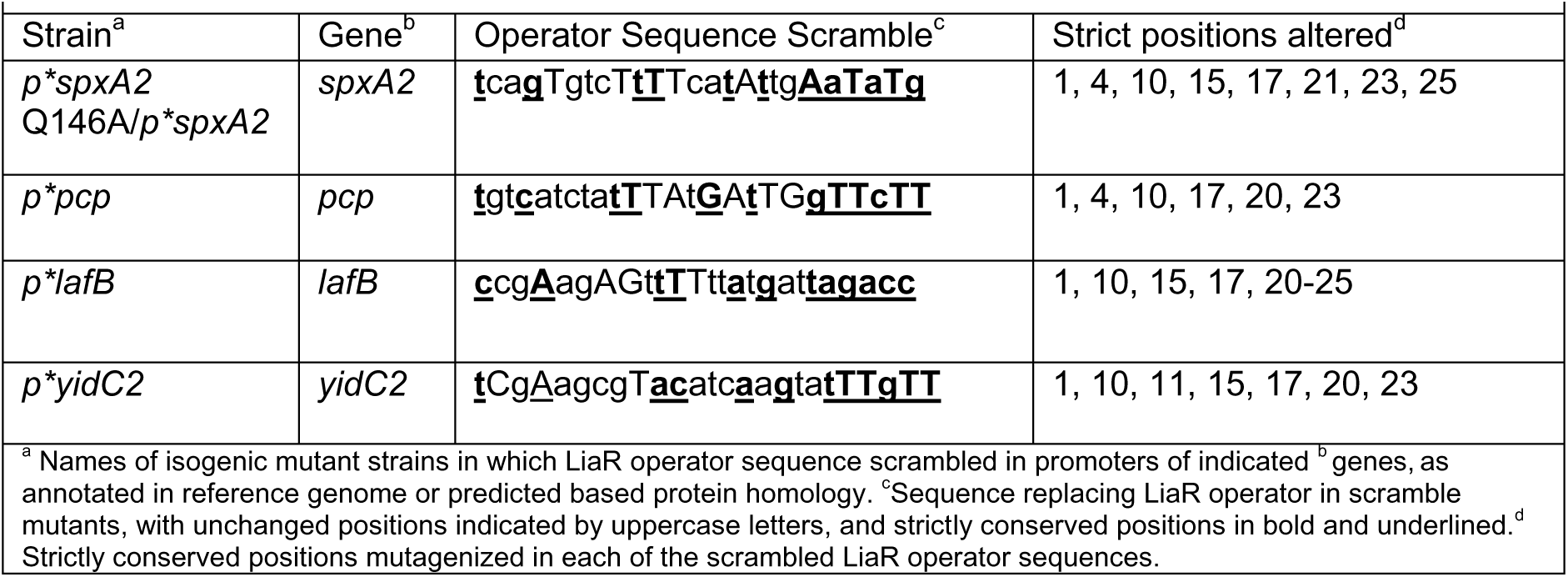
LiaR operator scramble mutations.

To further confirm that the observed effects on LiaR target transcription resulted from abrogation of operator interaction with phosphorylated LiaR, we mutagenized the *spxA2* promoter to scramble the LiaR binding motif in the LiaS-Q146A background (Q146A/p\**spxA2*; Table 3). Using qRT-PCR we determined levels of *spxA2* transcript were significantly lower in the p\**spxA2* relative to the Q146A parental strain, as well as relative to a wild-type strain (*P*<0.05; Supplemental Figure S2). Furthermore, we detected statistically significant changes in transcript levels of virulence genes *mac-1* and *grab* (*P*<0.05), identified by our RNA-seq data as differentially expressed in the R135G transcriptome (**Figure 1D**) and detected in our previous studies of Δ*liaF* and Δ*liaR* transcriptomes (M. A. Sanson et al. 2021). Additionally, differences in *spxA2* transcript abundance in the MGAS10870 wild-type strain and isogenic p\**spxA2*, Q146A and Q146A/p\**spxA2* strains directly correlated with levels of SpxA2 protein, as determined by Western blot analysis of GAS culture lysates (Supplemental Figure S3). Overall, our data indicate that phosphorylated LiaR directly controls transcription of *spxA2*, *lafB*, *pcp* and *yidC2* via a conserved operator in response to cell envelope stress. The association of transcriptional changes suggest that LiaR-mediated gene regulation further alters virulence gene expression.

### The LiaR-dependent regulon contributes to GAS antimicrobial susceptibility and host stress tolerance

Previously, we showed a reduced ability of the LiaS-R135G carrier mutant (Flores et al. 2015) and the Δ*liaR* deletion strain (M. A. Sanson et al. 2021) to grow *ex vivo* in whole human blood. We hypothesized that decreased survival in blood was a consequence of diminished capacity to tolerate cell envelope stress. To begin testing this hypothesis, we assessed the growth of LiaR phosphorylation (e.g., LiaS-Q146A and LiaS-D142A) and operator mutant strains (e.g., p\**spxA2*) in the presence of cell envelope-targeting antimicrobials known to activate LiaFSR (Lin et al. 2020). Minimum inhibitory concentrations (MICs) of bacitracin, vancomycin, nisin, polymyxin B, and daptomycin for mutant strains of interest showed relatively modest differences compared to the wildtype (MGAS10870) (Table 4). Bacitracin and polymyxin B exhibited the largest differences in MIC across multiple mutants relative to MGAS10870 and was most pronounced in the Q146A/p\**spxA2* strain.

**Table 4.** Antimicrobial minimum inhibitory concentration (MIC, µg/ml)

| **TABLE 4.** Antimicrobial minimum inhibitory concentration (MIC, µg/ml) |  |  |  |  |  |  |
| --- | --- | --- | --- | --- | --- | --- |
| **LiaR-P status** | **Strain** | **Bacitracin** | **Vancomycin** | **Nisin** | **Polymyxin B** | **Daptomycin** |
| (+) | *MGAS10870* | 2 | 0.25 | 24 | 20 | 3.2 |
| (--) | Δ*liaR* | **0.5** | 0.25 | **18** | **10** | **1.6** |
| (-) | R135G | **0.5** | 0.5 | 24 | **10** | **1.6** |
| (-) | D142A | **0.5** | 0.5 | 24 | **10** | **1.6** |
| (++) | Q146A | 0.75 | 0.5 | 48 | **10** | 3.2 |
| (++) | Q146A/*p*<sup>*</sup>*spxA2* | **0.25** | 0.25 | 24 | **5** | **1.6** |
| (+) | *p*<sup>*</sup>*spxA2*<sup>WT</sup> | **0.5** | 0.25 | 24 | **10** | **1.6** |
| (+) | *p*<sup>*</sup>*pcp* | **0.5** | 0.5 | 48 | **10** | 3.2 |
| (+) | *p*<sup>*</sup>*lafB* | **0.5** | 0.5 | 48 | **10** | 3.2 |
| (+) | *p*<sup>*</sup>*yidC2* | **0.5** | 0.5 | 24 | 20 | 3.2 |

We further assessed antimicrobial susceptibility by measuring the bactericidal activity of bacitracin and polymyxin B using virtual colony counts (CFUv; see Methods) (Ericksen et al. 2005). Strains with reduced LiaR phosphorylation (LiaR [-]) showed a significant reduction in survival as measured by CFUv following 90-minute exposure to bacitracin (*P*<0.05, **Figure 3A**) and polymyxin B (*P*<0.05, **Figure 3B**). Disruption of the LiaR operator of *spxA2* in both wild-type and Q146A backgrounds also resulted in a significant reduction in CFUv following exposure to bacitracin and polymyxin B, similar to the decreased survival of a Δ*liaR* mutant. In contrast, bacitracin and polymyxin B bactericidal activity on the Q146A strain was similar to that observed in the wild-type strain. LiaR operator sequence disruption in *pcp*, *lafB*, and *yidC2* promoters did not significantly alter bactericidal activity of the tested antimicrobials relative to an isogenic wildtype strain, except for the p\**yidC2* strain in polymyxin B (**Figure 3B**). Thus, these data indicate that decreased SpxA2 levels through the loss of LiaR regulation are the principal contributor to antimicrobial-mediated cell envelope stress.

**Figure 3.**
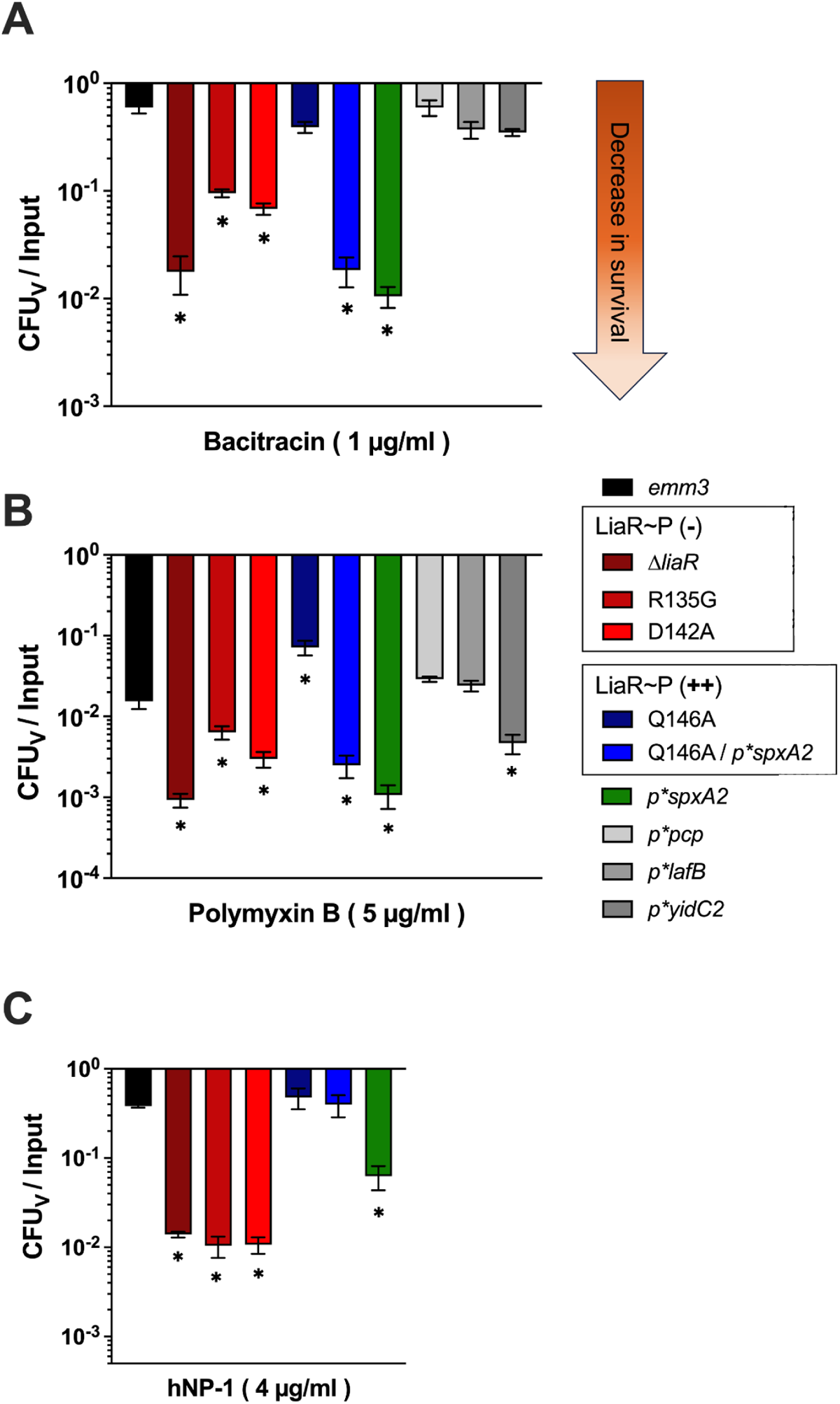
Disruption of LiaR phosphorylation and of LiaR regulon control alter the GAS response to AMPs. Ratio of surviving virtual colony counts (CFU_V_) to inoculum CFU counts (Input) of GAS suspensions (2-5 x 10^5^ CFU total) in Na-Phosphate buffer (10mM, pH 7.4) incubated for 90 minutes in (**A**) bacitracin, (**B**) polymyxin B and (**C**) hNP-1 indicates increased susceptibility to AMP bactericidal activity in strains lacking LiaR phosphorylation. The status of LiaR phosphorylation in tested strains is indicated the figure legends ([LiaR∼P (-)] reduced LiaR phosphorylation, [LiaR∼P (++)]: constitutive LiaR phosphorylation). Data is presented as the mean value with standard error of the mean (SEM). Significance determined by a t-test (*,P < 0.05).

Inasmuch as GAS is a human-specific pathogen that responds to human molecules, we next performed the same CFUv determinations following exposure to hNP-1 – the host AMP known to activate the LiaFSR system (Lin et al. 2020). Strains in which LiaR was absent or phosphorylation was reduced showed a significant decrease in CFUv following exposure to hNP-1 (*P*<0.05, **Figure 3C**). Likewise, similar to the observation with bacitracin and polymyxin B exposure, disruption of the LiaR operator sequence of *spxA2* in the wildtype background resulted in significantly decreased survival (*P*<0.05). On the other hand, the Q146A mutant (high LiaR phosphorylation) showed no significant change in survival following exposure to hNP-1. Disruption of the LiaR operator in *spxA2* in the Q146A background (Q146A/p\**spxA2*) also did not alter GAS survival. These results indicate that LiaR phosphorylation and induction of *spxA2* transcription contribute to tolerance of hNP-1, but constitutive phosphorylation of LiaR in the Q146A background promotes hNP-1 tolerance independently of induction of *spxA2* transcription.

In previous studies, we showed that targeted gene deletion resulting in disruption of LiaFSR activity led to decreased survival in human blood *ex vivo* (i.e. Δ*liaF*, Δ*liaR*) (M. A. Sanson et al. 2021). Therefore, we hypothesized that, alteration of LiaR phosphorylation is responsible for the survival defect in human blood. To test this hypothesis, first we assessed the survival in whole human blood of our LiaR phosphorylation mutant strains. Both LiaR hyperphosphorylation (Q146A) and unphosphorylated LiaR (R135G, D142A) mutants showed significantly decreased survival in whole human blood (*P*<0.05, **Figure 4A**). This result suggested that alteration of wild-type LiaR phosphorylation patterns and the associated dysregulation of LiaR targets have detrimental effects on GAS survival. To further examine the contribution of LiaR targets, we assessed the survival in whole human blood of the scrambled

**Figure 4.**
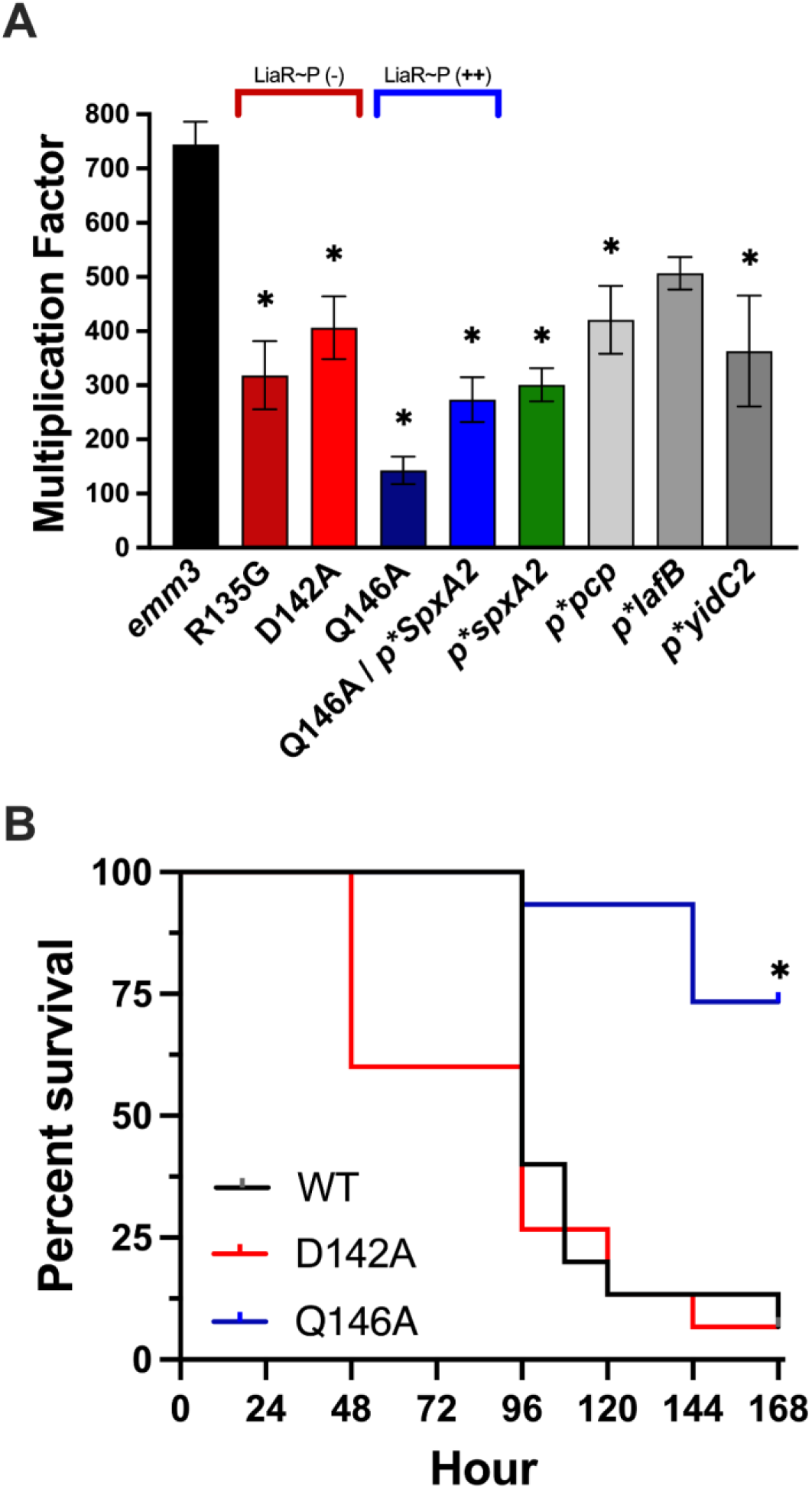
Disruption of LiaR phosphorylation and of LiaR regulon control negatively affect GAS virulence. (**A**) Survival in whole human blood (3 hours, 37°C) of strains in which LiaR phosphorylation is reduced [LiaR∼P(-)], LiaR is constitutively phosphorylated [LiaR∼P(++)], or the operator of LiaR regulon targets is mutagenized (Q146A/p\**spxA2*, p\**spxA2*^WT^, p\**pcp*, p\**lafB*, p\**yidC2*). The mean multiplication factor of GAS in whole human blood and SEM are shown, calculated from biological replicates in blood from 3 donors, performed in quadruplicate. Significance was determined by a t-test (*, *P* < 0.05). (**B**) Kaplan-Meier survival of mice (n = 15 per strain) infected intramuscularly with 2 x 10^7^ CFU D142A, Q146A, or MGAS10870 (WT). *P-*values were determined by log rank.

LiaR operator mutant strains. The mutagenized LiaR operator mutants showed significantly decreased survival in whole human blood to varying degrees (*P*<0.05, **Figure 4A**). The greatest differences relative to the wild-type strain were observed in the Q146A, Q146A/p\**spxA2* and p\**spxA2* strains, indicating that dysregulation of SpxA2 expression and LiaR dephosphorylation have the greatest influence on LiaFSR-dependent GAS fitness in whole blood. We further tested the effect of altered LiaR phosphorylation on GAS virulence using a mouse model of intramuscular GAS infection. Mice were infected intramuscularly with 1x10^7^ CFU MGAS10870 (wild type), Q146A (LiaR∼P++), or D142A (LiaR∼P-) isogenic mutant strains. In mice monitored over 7 days post-infection for signs of near mortality, the Q146A strain displayed significantly attenuated virulence (73% survival; *P*<0.05) relative to the isogenic parent MGAS10870 strain (7% survival) (**Figure 4B**). Conversely, mice infected with the D142A strain displayed no difference in survival relative to mice infected with the wild-type MGAS10870 strain, similar to what was observed previously in the same mouse model of infection with the R135G mutant (Flores et al. 2017).

## Discussion

Our study adds to our understanding of LiaFSR in Gram-positive bacteria and further elucidates LiaFSR function and its contribution to the GAS response to AMPs. Our data indicate that LiaR phosphorylation by LiaS in response to AMP-mediated cell envelope stress results in direct induction of a restricted regulon via a conserved operator. In addition to the previously identified transcriptional regulator *spxA2*, our experimental data identified four targets of direct regulation by phosphorylated LiaR: a PspC-domain containing protein (*pcp,* SpyM3_0411), a putative sugar transferase associated with lipoteichoic acid biosynthesis (*lafB*, SpyM3_0363) and a membrane protein insertase (*yidC2*, SpyM3_0256). Mutation of the LiaR operator resulted in dysregulation of phosphorylated LiaR targets and led to reduced GAS fitness in an *ex vivo* model of host infection. However, reduced LiaR phosphorylation did not attenuate virulence *in vivo*, whereas constitutive LiaR phosphorylation significantly attenuated virulence. Finally, the dysregulation of LiaR-dependent gene expression affected phenotypes associated with cell envelope homeostasis in GAS, particularly susceptibility to AMPs.

Dysregulation of SpxA2 expression had the largest effect on the GAS response to antimicrobials and AMPs indicating a key role in the GAS response to cell envelope stress. This observation is likely attributable to the broader effect of SpxA2 on global gene expression as a transcriptional regulator. Unique to and conserved across Gram-positive bacterial species, Spx proteins function as regulators of global gene expression controlling responses to oxidative stresses [for a comprehensive review, see (Rojas-Tapias and Helmann 2019)]. Characterization in *Bacillus subtilis* has revealed Spx proteins bind the α-C-terminal domain (CTD) of RNA polymerase (RNAP) to influence DNA, transcription factor, and σ-factor-binding of RNAP (Nakano et al. 2003; Zuber 2004). Through this mechanism, Spx proteins influence expression across the *B. subtilis* genome, regulating genes involved in the oxidative stress response, metabolism and transmembrane transport (Rochat et al. 2012). GAS encode two Spx paralogs (SpxA1 and SpxA2), which previous research has established have distinct effects on virulence, oxidative stress and antimicrobial tolerance (Port et al. 2017). Specifically, Port et. al. showed that loss of SpxA2 was associated with increased susceptibility to polymyxin B and oxidative stress. Disruption of the LiaR operator of *spxA2* reduced GAS survival to challenge by hNP-1, the major AMP produced by human neutrophils. Combined with the specific response of LiaFSR to α-defensins such as hNP-1 (Lin et al. 2020; M. A. Sanson et al. 2021), our data support a critical role of LiaR-mediated SpxA2 activity in GAS pathogenesis. GAS responses to host AMP activity are incompletely understood and prior work demonstrates activation of the TCS encoded by *ihk/irr* following exposure to neutrophil α-granules (Voyich et al. 2004) known to be rich in hNP-1 (Lehrer, Lichtenstein, and Ganz 1993). The mechanism by which LiaR (and by extension SpxA2) influences regulatory signaling and cross talk with other GAS TCS such as Ihk/Irr is unknown and an active area of investigation in the GAS response to human AMPs.

The current study also defines the mechanism by which the carrier mutation (LiaS-R135G) alters gene expression leading to what we termed a “carrier phenotype” defined by decreased virulence and increased ability to adhere/persist on epithelial surfaces (Flores et al. 2015; 2017). Both the LiaS-D142A and LiaS-R135G isogenic mutants dramatically reduce LiaR phosphorylation even in the presence of known LiaFSR activators. The inability to phosphorylate LiaR severely hampers the response to antimicrobial or AMP-induced cell envelope stress in large part through reduced *spxA2* gene expression. Paradoxically, our work and that of Port et. Al. (Port et al. 2017) showed that loss of SpxA2 enhanced GAS virulence but was dependent upon SpeB (M. A. Sanson et al. 2021). Given the pleiotropic regulatory effects of Spx proteins in other Gram-positive bacteria it remains highly likely that factors beyond SpeB contribute to the GAS LiaFSR/SpxA2 response during human infection.

Despite significant increases in virulence gene expression, constitutive phosphorylation of LiaR through loss of the *liaF* gene or through the LiaS-Q146A mutation results in decreased virulence *in vivo*. The contradictory effects of constitutive LiaR phosphorylation may be due, at least in part, to the disruption of anionic lipid localization in the GAS cytoplasmic membrane. In *Enterococcus faecalis*, reduced daptomycin susceptibility was associated with redistribution of cardiolipin-rich membrane microdomains away from the bacterial division septa in a manner dependent on LiaF mutation (Tran et al. 2013). Recent research further suggests that LiaFSR-dependent regulation of cardiolipin synthase expression is the mechanism responsible for this daptomycin resistance phenotype (Nguyen et al. 2023). Given the known association of the ExPortal with peptidoglycan synthesis (Vega, Port, and Caparon 2013), it is possible that anionic lipid redistribution associated with constitutive LiaR phosphorylation reduced AMP killing by disrupting interaction of the tested AMPs with their targets of action (e.g., bacitracin and hNP-1 interaction with lipid II). Concomitantly, the importance of ExPortal integrity to virulence factor secretion (J. W. Rosch and Caparon 2005; Vega and Caparon 2012) and pathogenesis (M. A. Sanson et al. 2021) would suggest that there is a fitness cost to constitutive LiaFSR activation. At this moment, whether constitutive LiaR phosphorylation leads directly to anionic lipid redistribution is unknown and the subject of future investigations.

In the current study, disruption of LiaR-dependent differential expression of *pcp* and *yidc2* by mutagenesis of the LiaR operator reduced GAS fitness *ex vivo*. In *Streptococcus mutans*, direct regulation of a PspC domain-containing protein (PCP) by LiaR was found to contribute to virulence phenotypes (biofilm formation, extracellular DNA release, platelet adhesion) in a model of infective endocarditis (Jung et al. 2021). The PspC domain corresponds to the protein of the same name that is an integral part of the phage shock protein (Psp) stress response system characterized in *Escherichia coli* and *Yersinia enterocolitica* (Joly et al. 2010). In the latter, PspC prevents toxicity of mislocalized secretins and forms an integral inner membrane complex with PspB that activates the Psp response (Maxson and Darwin 2006). Furthermore, a *pspC*-lacking strain of *Y*. *enterocolitica* shows attenuated virulence in a mouse infection model and PspC was required for bacterial growth under expression of the Type III secretion system (Darwin and Miller 2001). The function of Pcp in GAS and its contribution to pathogenesis is currently unknown.

In *S. mutans,* deletion of *yidC2* attenuates virulence *in vivo* (Palmer et al. 2012), and its loss is associated with aberrant cell wall synthesis, a reduction in biofilm biomass and pronounced defects in the spatial organization of the exopolysaccharide matrix (Palmer et al. 2019). Additionally, YidC2 appears to work in concert with the signal recognition particle (SRP) pathway in *S. mutans*, guiding integration into the cytoplasmic membrane of substrate proteins bound to the SRP receptor FtsY (Lara Vasquez et al. 2021). Previous work in GAS has shown that the SRP pathway is required for virulence *in vivo* and contributes to the secretion of Nga, SLO and SpeB virulence factors (J. W. Rosch et al. 2008). LiaR-dependent regulation of YidC2 expression may thus potentially contribute to the virulence response of GAS to host AMP-induced envelope stress. Future work aims to elucidate the contribution of LiaR-regulated targets to GAS virulence and their mechanism of action.

In summary, our studies demonstrate that the LiaFSR system of GAS primarily exerts its regulatory response to antimicrobial activity via LiaR-dependent control of SpxA2 expression. Our data highlight the species-specific nature of the LiaR-dependent bacterial transcriptomes involved in the response to cell envelope-targeting AMPs. In other Gram-positive pathogens studied (e.g. *E. faecalis, Enterococcus faecium, B. subtilis, Listeria monocytogenes, S. agalactiae*), LiaFSR activation targets a variety of factors distinct from those of GAS. Such LiaR-regulated factors include additional LiaR operon genes (e.g. LiaIGH) (Jordan et al. 2006), secreted signaling peptides (LiaXYZ) (Axell-House et al. 2024), ABC transporters (Fritsch et al. 2011; Chilambi et al. 2020), and factors involved in lipid membrane remodeling (Chilambi et al. 2020; Nguyen et al. 2023). Our studies further demonstrate that the LiaFSR systems of Gram-positive pathogens, while highly conserved, act via clearly distinct mechanisms likely reflective of the unique environmental niche each species occupies.

## Methods

### Bacterial strain and culture conditions

The strains used in this study are listed in Supplemental Table S3. GAS strains were grown in Todd-Hewitt broth containing 0.2% (wt/vol) yeast extract (THY broth; Difco Laboratories), on THY agar, or on Trypticase soy agar (TSA II) containing 5% sheep blood agar (SB; Becton, Dickinson). For all *in vitro* assays, unless otherwise indicated, cultures were grown overnight without shaking in THY broth at 37°C with 5% CO_2_ and used to inoculate fresh, prewarmed THY broth for growth to the culture density required.

### Generation of isogenic mutant strains

The primers used in this study are listed in Supplemental Table S4. Isogenic mutants lacking LiaF and LiaR proteins, as well as the LiaS-R135G substitution mutant lacking in LiaR phosphorylation were generated in previous studies (Lin et al. 2020; Flores et al. 2015). Isogenic mutants in which the pattern of LiaR phosphorylation was altered were generated in the *emm3* strain background (MGAS10870) by the in-frame replacement of nucleotides in the *liaS* (*SpyM3_1367*) coding sequence resulting in the following residue substitutions: glutamine to alanine at position 146 (Q146A), aspartate to alanine at position 142 (D142A). Mutations were generated as detailed in (Flores et al. 2015), with modifications. Briefly, complementary primers encoding single nucleotide substitutions at LiaS amino acid positions 142 and 146 were used to amplify flanking regions of homology in the *liaFSR* coding region (1000bp in the 5’ and 3’ direction) and subsequently ligated into the *E. coli*-Gram positive shuttle vector pJL1055 (Gryllos et al. 2008; Li et al. 1997) using *BamH*I and *Xho*I restriction sites. Electrocompetent cells of MGAS10870 were transformed with these plasmids and allelic replacement carried out as previously described (Flores et al. 2013; Carroll et al. 2011).

Mutants in which the LiaR operator sequence was scrambled were generated by in-frame allelic replacement in MGAS10870 background with the sequences specified in Table 2 and Table S3, using a counterselection approach employing a levansucrase (*sacB*) marker as detailed previously (Vega et al. 2022) with modifications. Briefly, overlap extension PCR was used to generate scrambled operator sequences flanked by 200-500bp homology arms for each LiaR target promoter for insertion into the pMAS39 vector (pJL1055 backbone containing *sacB* cassette) using *BamH*I and *Xho*I restriction sites. The obtained plasmids were introduced into electrocompetent GAS generated as previously described (Sitkiewicz and Musser 2006), substituting 0.5M polyethylene glycol (PEG) for sucrose in the growth and recovery medium to allow for counterselection, and transformants were screened for vector integration on THY agar supplemented with chloramphenicol (10µg/ml). Recombinants were then selected on THY agar supplemented with 150mM sucrose, and recombinants encoding scrambled LiaR operator sequences identified by colony PCR using primers specific to both the scrambled and native operator sequences. Whole-genome sequencing of recombinant strains confirmed the lack of spurious mutations.

### RNA-sequencing (RNA-seq) and qRT-PCR analysis

Transcriptional analyses were performed according to previously described protocols (M. Sanson and Flores 2020) with modifications. Briefly, For RNA-seq analysis of GAS strains exhibiting high (Q146A) and low levels of LiaR phosphorylation (R135G), GAS cultures were grown in THY (biological quadruplicate) to exponential phase (OD600 0.9-1.0), and cells harvested by centrifugation following the addition of RNAprotect bacteria reagent (Qiagen). RNA was isolated and purified with a RNeasy mini kit (Qiagen) according to the protocol for Gram-positive bacteria. The quantity and quality of RNA were determined using NanoDrop (ThermoFisher) and an Agilent 4200 TapeStation system (Agilent Technologies). RNA purification, rRNA depletion and development of adapter-tagged cDNA libraries were performed using Script-Seq Complete Kit (Bacteria) [Epicentre (Illumina)]. cDNA libraries were run on a NextSeq 2000 and an average of ∼62 million (range 48-77 million) 150-bp paired-end reads were obtained per replicate sample. Following on-board adapter trimming and quality control, reads were subsequently mapped to the reference *emm3* genome (MGAS10870; Accession number CP067090.1) using CLC Genomics Workbench version 23 (Qiagen). Identification of significantly differentially expressed genes was determined as we have previously described (M. Sanson and Flores 2020). Genes for which the change in expression was >1.5 fold and *P-*value <0.05 following Bonferroni’s correction for multiple comparisons were considered to have a significant difference in expression.

RNA from GAS cultures for qRT-PCR analysis was isolated from cultures grown to exponential phase (OD600 0.5-0.6) and either exposed to bacitracin (1µg/ml final concentration) or left untreated for 1h (final OD600 ∼1.0), 37°C in 5% CO_2_ and purified with a RNeasy mini kit (Qiagen) according to the protocol for Gram-positive bacteria. The quantity and quality of RNA were determined as indicated above. TaqMan (Applied Biosystems) quantitative real-time PCR (qRT-PCR) of cDNA generated using SuperScript III reverse transcriptase (Invitrogen) was performed with an CFX96 TouchTM Real-Time PCR detection system (Bio-Rad) according to the manufacturer’s instructions. The TaqMan primers and probes used in the analyses are listed in Table S4 (Supplemental Data). Transcript levels of genes were calculated by relative quantification using the ΔΔCT method as described (Livak and Schmittgen 2001), with an internal reference gene *tufA* (*SpyM3_0432*) as the normalizing gene. All reactions were performed in triplicate using RNA purified from four biological replicates.

### Chromatin immunoprecipitation sequencing (ChIP-seq) analysis

Three biological replicates of strains MGAS10870 (low LiaR∼P), LiaS-Q146A (high LiaR∼P), and Δ*liaR* (control) were chromatin immunoprecipitated using a previously generated anti-LiaR antibody (Lin et al. 2020). Briefly, strains were grown to mid-logarithmic phase (OD600 ∼0.4) in 40 mL THY, proteins crosslinked to DNA with 1% formaldehyde, cells harvested by centrifugation, flash-frozen, and stored at -80°C until processed. Processing of replicate samples for ChIP-seq was performed as previously described (Horstmann et al. 2023; DebRoy et al. 2021). High-throughput sequencing on prepared sample libraries was performed by Genewiz (Azenta Life Sciences) to generate a minimum of 50M 150-bp paired reads per replicate/input sample. Analysis was performed as previously described (Horstmann et al. 2023; DebRoy et al. 2021) using CLC Genomics Workbench (v 23) Transcription Factor ChIP-seq module.

### Antimicrobial susceptibility measurements

For antimicrobial susceptibility measurements via virtual colony counts (CFU_V_), we employed a previously published approach (Ericksen et al. 2005) with modifications. Seed cultures for susceptibility testing were grown from single colonies from TSA-II/SB plates in 10 ml of THY. After overnight growth, 500 μl of culture was added to 10 ml of THY and grown to an OD_650_ of 0.45 to 0.55 to obtain ∼10^8^ CFU/ml of GAS. A 20-fold dilution of this culture in sodium phosphate buffer (10mM, pH 7.4) was used to generate a cell suspension of ∼5x10^6^ CFU/ml.

Antimicrobials (AMPs) were diluted to 2X final desired concentration in 10mM sodium phosphate buffer (pH 7.4). AMP solutions were then aliquoted (50µl) in triplicate per strain and concentration tested in a 96-well plate format (AMP-treatment plate). GAS cell suspension (50µl) was added to each AMP-containing well and to sodium phosphate buffer-only wells (untreated samples) to obtain ∼5x10^5^ CFU/well in a 100µl volume. The AMP-treatment plate was then incubated for 90 minutes (37°C, 5% CO_2_). Following incubation, 100µl of double-strength (2x) THY was added to each of the wells and mixed thoroughly. A 20µl aliquot from each well of the AMP-treatment plate was then transferred to 180µl of fresh, pre-warmed (37°C) THY in a clear bottom 96-well plate (Measurement plate) for outgrowth and optical density quantification. A 10-fold dilution series (10^4^ to 10^1^) of additional 20µl aliquots from each of the untreated sample wells was plated on TSA-II/SB and incubated overnight for CFU quantification to determine the starting concentration of GAS (*C*’_0_) treated. Optical density measurements (OD_600_) were performed every 5 minutes over 20 hours on a Synergy H1 (Biotek) microplate reader (chamber supplemented with 5% CO_2_, 37°C), with shaking for 30 seconds initially and then for 10 seconds at every 5 min time point thereafter. For generation of the calibration growth curves used for CFU_V_ interpolation, a 10-fold dilution series of the untreated sample wells (10^4^ to 10^1^ CFU/well) was made in the Measurement plate in a final 200 μl THY volume. Wells containing only THY medium were included as blanks for normalization.

Prior to AMP susceptibility testing of isogenic mutant strains, hNP-1 and polymyxin B concentrations for testing were identified by determining the LD_90_ concentrations (i.e. AMP amounts resulting in <10% survival) of the isogenic parental *emm3* strain (MGAS10870). To that end, a twofold dilution series of AMPs was tested with concentrations ranging from 12 to 2.0 μg/ml (hNP-1), and 10 to 0.83 µg/ml (polymyxin B; Supplemental Figure S4). The concentration of bacitracin chosen for isogenic mutant comparisons was determined from previous experimentation indicating consistently reproducible activation of the LiaFSR system upon GAS exposure to 1µg/ml bacitracin (M. A. Sanson et al. 2021; Lin et al. 2020). Each strain was tested four times (biological replicates) on separate days by starting with overnight cultures inoculated from different TSA-II/SB plates.

For analysis, the OD_600_ measurements from the 20-h plate reader run were normalized by subtracting the blank well readings from each well’s readings. Growth curves for each sample (i.e. strain at a given AMP concentration) tested were generated using the median values of the technical replicates (3) in which they were measured. The times at which each untreated sample growth curve crossed the threshold values of 0.01, 0.02, 0.05, 0.1, and 0.2 absorbance units were plotted against log(*C*’_0_) to generate a calibration curve that related the threshold times to *C*′_0_. Linear regression (*r*^2^) values for calibration curves used for interpolation ranged from 0.98 to 0.999 at a threshold ΔOD_600_ of 0.02 and similar slopes, *y* intercepts, and *r*^2^ values were obtained at all thresholds from 0.005 to 0.4. The times at which AMP-treatment growth curves crossed threshold absorbance unit values were interpolated into the calibration curve corresponding to each strain to obtain CFU_V_. Outlier CFU_V_ values were detected and eliminated using Prism (GraphPad) analysis (Q=5%). The survival ratio for each strain at each AMP concentration tested was calculated as the CFU_V_ AMP-treated/*C*′_0_.

For AMP minimum inhibitory concentration (MIC) determination, Clinical and Laboratory Standards Institute (CLSI) guidelines for MIC determination using broth microdilution in liquid Mueller Hinton broth (MHB) were followed (*M100 PERFORMANCE STANDARDS FOR ANTIMICROBIAL SUSCEPTIBILITY TESTING, 33RD EDITION,M100ED33* 2023).

### Survival in whole human blood

The ability of GAS to survive in whole human blood were conducted under a human subject protocol approved by the UTHealth Houston Committee for the Protection of Human Subjects (CPHS protocol #HSC-MS-17-0875). Experiments were carried out as described previously (Lancefield 1957). Briefly, blood samples from three healthy, nonimmune, adult donors (2 female and 1 male) were used as biological replicates, and assays were performed in quadruplicate (technical replicates). GAS inocula of ∼50 CFU were incubated for 3h, at 37°C with 300µl of donor blood with rotation end over end. Surviving CFU were measured by plating of serial dilutions on TSA II-SB (Becton, Dickinson).

### Mouse model of intramuscular infection

All animal experiments were conducted under a protocol approved by the UTHealth Houston Institutional Animal Care and Use Committee. CD1 mice (Charles River, 3 to 4 weeks of age, male and female) were inoculated in the right hind limb with 5 × 10^7^ CFU of the appropriate GAS strain in 50 μl of phosphate-buffered saline (PBS). For survival, infected mice (*n* = 15 per strain group) were observed and euthanized when they reached near mortality determined using predefined criteria.

### Statistical analysis

All statistical analyses were performed in Prism 10 (GraphPad Software, Inc.). Differences between mean values were tested using a two-tailed Student’s *t-*test, Mann-Whitney U (nonparametric data), and statistical significance set to a *P* value of <0.05 for all comparisons.

### Data availability

All RNA-seq and ChIP-seq data have been deposited in the NCBI GEO database under accession number GSEXXXXXX.

## Acknowledgments

This work was supported by funding provided by the National Institute of Allergy and Infectious Diseases R01 AI124216 (A.R.F.).

